# Stable yet Shifting: Early Toxin Dynamics in Typical and Atypical Clownfish–Anemone Symbioses

**DOI:** 10.64898/2026.05.26.727870

**Authors:** J. Macrander, A. Bennett, K. Statile, W. Rudd, C. Tolman, S. Kuklina, S. Burg, L. Whitton, G. Langford

## Abstract

Among venomous animals, cnidarians represent the oldest metazoan lineage in which venom production and a specialized delivery system are defining synapomorphies. Cnidarians also represent the only venomous lineage for which mutualistic symbioses have evolved resulting in scenarios where mutualistic symbionts may also be targets of their venom. The most iconic example of this relationship is the mutualism between clownfish and their venomous sea anemone hosts. To investigate how symbiont presence and establishment influence toxin gene expression, we used a comparative TagSeq and RNA-Seq approach to quantify venom gene dynamics during the first 48 hours of clownfish–anemone symbiosis establishment in five anemone species. Our taxonomic sampling included three typical hosting species (*Entacmaea quadricolor*, *Radianthus crispa*, and *Stichodactyla haddoni*), each representing distinct evolutionary lineages of clownfish hosts, and two atypical Caribbean species (*Condylactis gigantea* and *Stichodactyla helianthus*) that do not host clownfish in nature, but have reported to host within the aquarium trade. Tentacle samples were collected prior to hosting, approximately 12 hours after initial symbiont establishment, and again 48 hours after symbiosis establishment. Our analyses revealed that overall toxin assemblages remained relatively stable during the early establishment phase, with no significant changes in the most highly expressed toxin gene candidates. However, subtle transcript-level shifts occurred within multi-copy toxin gene families, including cytolytic actinoporins and Sea Anemone 8 (SA8)-like toxins. Notably, one *C. gigantea* actinoporin transcript exhibited a ∼600-fold increase in expression in a single individual, which coincided with two clownfish mortalities prior to successful association, which subsequently decreased after establishment. Comparative sequence alignments suggest that amino acid substitutions in this transcript may be functionally relevant to symbiosis intolerance, as the amino acid substitutions were unique to this transcript, and not found in any other previously described cytolytic actinoporin. Together, these findings reveal that early toxin gene expression in clownfish-hosting sea anemones is largely stable, yet subtly dynamic at the transcript level. This study provides the first comparative transcriptomic insights into the molecular processes shaping symbiosis establishment in clownfish–anemone mutualisms, offering a framework for understanding venom evolution in the context of co-evolutionary interactions.

**Highlights:**

- Comparative gene expression survey reveals relatively stable toxin assemblages throughout the first 48 hours of establishing clownfish-anemone symbiosis.
- Subtle shifts were observed among transcript variants in multi-gene copy variants, with potential implications for barriers to establishing symbiosis.
- Although toxin assemblages varied among species, sea anemone 8 (SA8) toxin-like transcripts were highly abundant in four of the focal taxa.
- This is the first comparative gene expression analysis investigating molecular processes surrounding symbiosis establishment between clownfish and sea anemones.
- These results provide insight into toxin dynamics surrounding the establishment of symbiosis, with particular insights into key evolutionary transitions resulting in symbiosis among atypical clownfish hosting species.

## Introduction

Cnidarians represent the earliest diverging metazoan lineage in which venom production and nematocyst-based delivery are defining synapomorphies (Ashwood et al., 2020; Daly et al., 2007; D’Ambra and Lauritano, 2020; Jouiaei et al., 2015a). They exhibit developmental and anatomically specific toxin gene expression profiles (Ashwood et al., 2021; Macrander et al., 2016, 2015; H. L. Smith et al., 2023a), modulation of venom transcription in response to environmental stressors (Sachkova et al., 2020), and express a diversity of bioactive compounds with biomedical potential (Liao et al., 2019; Mariottini and Pane, 2014; Merquiol et al., 2019; Turk and Kem, 2009). While these features are broadly shared among venomous taxa, cnidarians, particularly sea anemones (Actiniaria), are unique in forming close physical associations with species that would otherwise be susceptible to their venom. Several crustaceans, including *Periclimenes*, *Thor*, and *Ancylomenes* inhabit sea anemone tentacles as cleaner and commensal shrimps (Briones-Fourzán et al., 2012; Mascaró et al., 2012). Hermit crabs, such as *Pagurus bernhardus* and *Dardanus pedunculatus* attach *Calliactis* spp. sea anemones to their shells for protection (Ross, 1971). The boxer crabs (*Lybia sp.*) carry sea anemones in their claws as kleptoparasitic extensions to be used for defense and capturing food (Schnytzer et al., 2022). Even deep sea gastropods and their sea anemone associates benefit from these mutualistic relationships providing protection and more efficient foraging (Mercier et al., 2011). Beyond these invertebrate examples, the mutualism between clownfish and sea anemones remains the best studied example of this phenomenon (Fautin, 1991; Hoepner et al., 2022; Lubbock, 1981; Porat and Chadwick-Furman, 2004), where host sea anemones harbor fishes and crustaceans that, under other circumstances, would represent natural targets for their venom (Giese et al., 1996; Mebs, 2009).

In contrast to their invertebrate counterparts, the mutualisms that occur between anemonefish and sea anemones encompass dozens of species across multiple lineages. This mutualism can be found widely across the tropical Indo-West Pacific, comprising 28 species of anemonefish and 10 sea anemone species (Fautin, 1991; McCord et al., 2021; Titus et al., 2024). These partnerships vary along a continuum from highly specialized to broadly generalist associations (Fautin, 1991; Hoepner et al., 2022; Litsios et al., 2014b, 2012; Marcionetti et al., 2019). Phylogenetic reconstructions among the anemonefish have suggested that the establishment of this symbiosis served as a catalyst for rapid recent speciation (Litsios et al., 2014a), while the capacity to host anemonefish has evolved independently at least three times within distinct lineages of sea anemones (Kashimoto et al., 2022; Titus et al., 2019), varying significantly in shape, color, and general morphology across their distribution (Titus et al., 2024). Once established, anemonefish are able to avoid being stung through a combination of behavioral and physiological adaptations (Balamurugan et al., 2014; Elliott et al., 1994; Nguyen et al., 2024), with anemonefish being susceptible to the venom if acclimation does not occur (Elliott and Mariscal, 1997; Mebs, 2009). Although the natural associations among species specific anemonefish and sea anemone have long been a focus of how mutualisms evolve in nature (Arvedlund et al., 1999; Chiodo et al., 2024; da Silva and Nedosyko, 2016), there are examples of symbioses being established outside of their typical host species without the anemonefish succumbing to the venom (Nguyen et al., 2024). Very little is known about how symbiotic relationships within clownfishes have co-evolved with venomous hosts over long-term evolutionary timescales, focusing on potential adaptive behaviors, venom inhibitors, and chemical signaling between the symbionts (Verde et al., 2015)

The coevolution of anemonefish and venomous host has prompted multiple studies characterizing the toxin repertoires of clownfish-hosting sea anemones through combined transcriptomic and proteomic approaches (Chiodo et al., 2024; Delgado et al., 2022; Hoepner et al., 2024; Hua et al., 2024; Madio et al., 2017). Venom gene expression in these species have been examined across multiple tissues (Ashwood et al., 2021; Macrander et al., 2016), with recent long-read genome assemblies further expanding the genomic toolkit available for evaluating venom diversity and regulation (De Jode and Titus, 2025). In other sea anemones, retention of toxin gene copies through concerted evolution and dosage compensation mechanisms have been documented (Moran et al., 2008; Surm et al., 2019), enabling bursts of diversification (Jouiaei et al., 2015a; Smith et al., 2023b), a likely adaptive advantage for taxa engaged in dynamic mutualistic interactions. Environmental heterogeneity across a broad range of habitats occupied by sea anemones has further shaped venom composition (Sachkova et al., 2020; Sunagar et al., 2016), which is further diversified through sub-functionalization, with both positive and negative selection acting across tandemly duplicated toxin gene copies (Sachkova et al., 2019; Sunagar et al., 2016). The evolution of eukaryotic mutualistic associations have been linked to dramatic ecological shifts and drastic changes in genomic content and structure (Doebeli and Knowlton, 1998; Hurst, 2017). Among venomous taxa, however, the formation of close, non-predatory associations with potential venom targets appears to be unique to Cnidaria, representing an ecological innovation likely to have shaped the evolution and functional complexity of their venom systems.

The best characterized sea anemone toxins include neurotoxins that target sodium (NaTx) and potassium (KTx) channels, along with membrane-disrupting cytolysins of the actinoporin family (Anderluh and Maček, 2002; Jouiaei et al., 2015b; Madio et al., 2019; Prentis et al., 2018). Transcriptomic datasets consistently recover these toxin gene families across all sea anemones, although the dominant toxin types vary substantially among superfamilies (Smith et al., 2023b). Notably, the high overall expression levels of each toxin family are often driven by one or two transcripts rather than evenly distributed among paralogs. This suggests that selection pressures may favor specific gene copies within genomic regions expressing multiple copy number variants (CNVs). This pattern implies environmentally or developmentally contingent regulation, where alternative CNVs may be upregulated under shifting ecological conditions. Recent analyses of clownfish-hosting sea anemones reveal venom repertoires dominated by hemolytic and hemorrhagic toxins (Delgado et al., 2022). However, the functional roles of many of these remain poorly understood and their relative contributions to the overall toxin assemblage remain unknown. From a physiological perspective, toxin gene expression is metabolically expensive (Sachkova et al., 2020) and if sea anemones are like other venomous taxa, a diverse non-specific venom phenotype would benefit both passive prey capture and defense (Lyons et al., 2020; Pekár et al., 2018; Phuong et al., 2016). In clownfish-hosting species, symbiotic interactions may act as a selective filter shaping venom composition by suppressing expression of antagonistic toxins and favoring the retention or upregulation of gene copies that are neutral or beneficial to symbiosis.

For these reasons, the venom phenotypes of clownfish-hosting sea anemones represent complex traits likely shaped by ecological interactions, with major implications for venom gene expression dynamics, gene family evolution, and the retention of CNVs across both micro- and macro-evolutionary time scales. To determine whether initial establishment of clownfish mutualistic symbioses drives a shift in toxin gene assemblages, we monitored the formation of associations across five species of sea anemones. This included three typical hosting sea anemones, *Entacmaea quadricolor*, *Radianthus crispa*, and *Stichodactyla haddoni*, each representing one of the three clades of hosting sea anemones, as well as two atypical hosting sea anemone species, *S. helianthus* and *Condylactis gigantea*. Following acclimation, candidate toxin gene expression in tentacles from each sea anemone were quantified prior to the introduction of a generalist clownfish (*Amphiprion clarkii*) and subsequently sampled again after 12 and 48 hours post association. To our knowledge, this study provides the first characterization of toxin gene expression dynamics accompanying the onset of symbiosis between clownfish and their sea anemone hosts.

## Materials and Methods

### Animal care, experimental setup, and hosting observations

In total, five species of sea anemone were used to assess the impact of clownfish association on sea anemone gene expression in three typical clownfish hosting sea anemones (*E. quadricolor*, *R. crispa,* and *S. haddoni*) and two atypical hosting sea anemones (*S. helianthus* and *C. gigantea*). Animals were provided from a personal/home aquarium (clonal *E. quadricolor*) or local aquarium suppliers (*C. gigantea*, *S. haddoni*, *H. crispa* and *S. helianthus*), which acquired them from national or international distributors with no known geographic origin. Sea anemones were housed in a flow through system consisting of nine 10-gallon aquariums. Aquariums were maintained at a constant temperature of 25℃, 30-33ppt salinity, with lights cycling through 12:12 light:dark cycles. In total, 6-8 animals of each species were used, with a single animal in each aquarium. Additional aquariums were used to house animals in reserve or clownfish waiting for the exposure experiment. Each species was tested separately due to limits on the number of aquariums available for the flow through system. At the start of the experiment a single clownfish (*Amphiprion clarkii*) was added to half of the aquariums containing sea anemones (3-4) following a minimum one-week acclimation period. The Clarkii clownfish were used as they are commonly reported as being broad generalists in symbiosis with a variety of different sea anemone species (Litsios et al., 2014a), including reported observations of atypical hosting species in aquaria. When possible, naive clownfish (those with no prior exposure to sea anemones) were used to minimize behavioral variance. However, due to limited animal availability and the prioritization of broad taxonomic sampling over replicate depth, the ‘naivety’ of the fish was treated as a descriptive observation rather than a controlled experimental variable, as the current sample sizes do not support a multi-factor statistical analysis of host experience. Prior to the start of the experiment clownfish and sea anemones were fed 1 mL of live *Artemia* napluii or frozen mysis shrimp three times per week. Water quality was measured along with cleanings and water changes were done on a weekly basis. Animal care guidelines were approved under IACUC protocols FSC-111621 and FSC-2023-04.

At the start of the experiment 3-6 tentacles were removed from each sea anemone using tweezers, placed in a 1.5 mL tube, and flash frozen in liquid nitrogen. Tweezers were then rinsed in deionized (DI) water and wiped clean for subsequent tentacle extraction. Clownfish were then randomly assigned to half of tanks containing sea anemones and observations were conducted to determine when clownfish hosting was established, with time-lapse cameras (TL2100, Dsoon, Shenzhen, China) to determine if establishment of symbiosis occurred overnight. Once the clownfish association was established tentacles were taken at approximately 12 hours post association and again approximately after 48 hours. For *E. quadricolor*, tentacles were sampled at 8 AM and 8 PM for the 48-hours sampling period to determine if circadian cycling influenced toxin gene expression. The exposure experiments were conducted for *E. quadricolor* and *C. gigantea* Spring 2021 and *H. crispa, S. helianthus*, and *S. haddoni* Summer 2023.

### RNA extraction and sequencing

Total RNA extractions for flash frozen tentacles were conducted using Purelink^TM^ kit (Invitrogen, Waltham, MA, United States) for *E. quadricolor* and *C. gigantea* and the TRIzol standard protocol (Invitrogen, Waltham, MA, United States) for *H. crispa, S. helianthus*, and *S. haddoni*. Once resuspended in RNAse free water all samples were quantified using a NanoDrop ND-1000 Spectrophotometer (ThermoFisher Scientific,Waltham, Massachusetts, United States) and samples stored at −80° C. Samples were then sent to Admera Health Biopharma Services, South Plainfield, NJ for sequencing. Once there, RNA quality and integrity was quantified on the Agilent TapeStation System (Agilent Technologies, Satna Clara, California, USA).

To assemble robust transcriptomes and optimize the number of samples for each species, we used a combination of previously published data along with TagSeq and RNA-Seq approaches. Based on RNA quality and integrity scores, we opted to sequence only the best three RNA samples for each time point for *E. quadricolor* and *C. gigantea* to serve as biological replicates. We sequenced the RNA using the TagSeq sequencing (1× 100 bp) approach (Lohman et al., 2016), with libraries constructed using the Lexogen QuantSeq kit (Lexogen, Vienna, Austria). The libraries were sequenced on the Illumina HiSeq 2500 with a minimum of four million paired reads per sample. As extensive paired end (2×150 bp) raw read data was already available for these species (BioProject: PRJEB21970) we opted to only conduct TagSeq sequencing in order to include biological replicates. Alternatively, we had to combine our biological replicates for *H. crispa, S. helianthus*, and *S. haddoni*, resulting in time-point specific RNA-Seq samples for each species. For these samples paired-end (2×150 bp) RNA-Seq libraries were constructed using the Illumina TruSeq Stranded mRNA Kit (San Diego, California, USA) and sequenced on the Illumina NovaSeq X, with a minimum of twenty million reads per sample. All sequencing was done by Admera Health Biopharma Services. Raw sequence data were then inspected using the program FastQC (Andrews, 2010) prior to downstream assembly or analyses. Raw reads for the TagSeq and RNA-Seq data were deposited on Genbank SRA (BioProject: PRJNA1469652).

### Transcriptome assembly, differential gene expression, and toxin gene identification

Using the program Trinity v2.13.2 (Haas et al., 2013) we assembled *de novo* transcriptomes for each species using either (1) publicly available raw reads for *E. quadricolor* (ERR2045166 -ERR2045171) and *C. gigantea* (ERR2045160 -ERR2045165), (2) a combination of publicly available raw reads for *R. crispa* (SRX19719912 -SRX19719918), *S. helianthus* (SRR7126073), and *S. haddoni* (SRX19719919 -SRX19719923) and our new RNA-Seq data or (3) a *de novo* assembly using just our raw reads across all species (Table 1). Once assembled, overall transcriptome completeness was determined using BUSCO (Simão et al., 2015) using the eukaryota_odb10 database. To quantify transcript abundance, raw reads were mapped to their respective transcriptomes using kallisto v0.48.0 (Bray et al., 2016). For whole-transcriptome RNA-Seq libraries (*R. crispa*, *S. haddoni*, and *S. helianthus*), abundance was quantified using Transcripts Per Million (TPM) to account for gene length bias. In contrast, the libraries for E. quadricolor and C. gigantea were quantified using Counts Per Million (CPM) as TagSeq generates reads independently of transcript length. This dual-normalization approach ensures that expression levels are accurately represented for each sequencing technology. We also mapped raw reads from the respective BioSamples for *E. quadricolor* and *C. gigantea* to further assess toxin diversity among individuals as our *E. quadricolor* sea anemones were known asexual propagules. Transcript variants inferred by Trinity were treated as putative isoforms, lacking a reference genome for all species, these sequences may represent distinct loci resulting from gene duplication events rather than alternative splicing. Toxin gene candidates were identified using the Tox-prot animal venom database limiting taxonomic sampling to Cnidaria (downloaded 4 June 2025). Assembled transcriptomes were screened for toxin candidates using tblastn from NCBI BLAST + v.2.1.2 with an e-value cutoff of 0.001. For species with biological replicates (*E. quadricolor* and *C. gigantea*) differential gene expression analyses across distinct time points during symbiont association were conducted using edgeR with a fold change of ≥4 and ≤0.05 p-value for significance (Robinson et al., 2010). Actinoporin toxin gene transcripts of interest were further evaluated by conducting sequence alignments using MAFFT (Katoh and Standley, 2013) to identify unique amino acids among transcript variants.

**Table 1.** Species specific transcriptomes, mapping, and toxin identification results.

| Species | Hosting | Sequencing | SRA | transcripts | BUSCO | N50 | Toxin Transcripts | pseudoalign % | DEG # |
| --- | --- | --- | --- | --- | --- | --- | --- | --- | --- |
| *E. quadricolor* | Typical | TagSeq<br>(1 x 100 bp) | ERR2045166 -<br>ERR2045171 | **442,451**<br>123,481 | **99.2%**<br>7.9% | **1,011**<br>339 | **1,195** | **56.9% - 74.1%**<br>24.0% - 42.9% | **715** |
| *C. gigantea* | Atypical | TagSeq<br>(1 x 100 bp) | ERR2045160 -<br>ERR2045165 | **244,876**<br>150,400 | **98.5%**<br>12.2% | **1,625**<br>411 | **1,034** | **62.9% - 71.7%**<br>29.9% - 55.1% | **251** |
| *S. haddoni* | Typical | RNA-Seq<br>(2 x 150 bp) | SRX19719919 -<br>SRX19719923 | **734,526**<br>562,579 | **95.7%**<br>61.5% | **1,043**<br>656 | **1,399** | **40.9% - 48.5%**<br>42.0% - 50.0% | NA |
| *R. crispa* | Typical | RNA-Seq<br>(2 x 150 bp) | SRX19719912 -<br>SRX19719918 | **605,746**<br>233,683 | **98.8%**<br>30.1% | **1,455**<br>453 | **1,355** | **30.9% - 50.9%**<br>30.6% - 54.1% | NA |
| *S. helianthus* | Atypical | RNA-Seq<br>(2 x 150 bp) | SRR7126073 | **735,730**<br>327,237 | **94.5%**<br>47.8% | **1,039**<br>565 | **1,392** | **37.4% - 40.5%**<br>48.2% - 55.3% | NA |

Bold items represent combined assemblies that were used as the focal analysis of this study.

## Results

### Association establishment and atypical acclimation results

For all species, associations took between 24 hours and four weeks to establish symbiotic associations among individuals. The typical hosting species (*E. quadricolor*, *R. crispa*, and *S. haddoni*) associated more quickly, with slightly faster associations being established if the *A. clarkii* clownfish had previously hosted with another sea anemone (Typically 24 −48 hours). Associations among the atypical hosting sea anemones were longer on average, however, one of the *C. gigantea* was associated in less than 24 hours after being introduced. For the others, after three weeks of no associations taking place between *C. gigantea* and *A. clarkii* we moved them to floating 500 ml containers with holes for water flow. Once in the container a second association was established within 48 hours. A single individual of *C. gigantea* ultimately took four weeks to establish a symbiotic association with clownfish due to apparent envenomation and ultimately the mortality of two *A. clarkii* clownfish (one naive, and one previously associated). For the atypical hosting sea anemone *S. helianthus* associations never took place for a smaller *A. clarkii* even after placing into floating 500 mL containers. Ultimately, a single large *A. clarkii* that readily hosted in other species began associating with *S. helianthus* allowing us to get tentacles for association across three individuals for three time points.

### Transcriptome assemblies, toxin identification, and mapping

The number of assembled transcripts varied significantly across species, ranging from 244,876 −734,526, with the largest number of transcripts recovered for *S. haddoni*. The de novo assembly of just our raw reads was done to evaluate overall completeness to prevent population specific genetic differences that could result in poor mapping or incomplete assemblies based on BUSCO scores. Ultimately, the assemblies based on either publicly available RNA-Seq data (*E. quadricolor* and *C. gigantea*) or a combination of publicly available and new sequence data for *R. crispa*, *S. haddoni*, and *S. helianthus* had the highest completeness with BUSCO scores greater than 94.5% across all focal taxa (Table 1) and used for downstream toxin analysis. As overall toxin assemblage for the clownfish hosting species was recently described (Delgado et al., 2022) our goal was to quantify expression of candidates, rather than categorize and sort, toxin assemblages found within each species. Using similar approaches for preliminary identification our initial BLAST hits recovered 1,034 −1,355 toxin transcripts. To quantify the pseudoalignment percentage varied across each species, with notable improvements for *E. quadricolor* and *C. gigantea* compared to the *de novo*, but slightly lower percentages for *R. crispa*, *S. haddoni*, and *S. helianthus* when compared to the de novo alone. Due to the higher BUSCO score for the combined, we opted to further evaluate toxin gene expression for the more complete transcriptomes to be consistent across species, but confirmed expression levels among toxin candidates among *R. crispa*, *S. haddoni*, and *S. helianthus* did not vary significantly regardless of assembly pipeline. For each species we chose to evaluate toxin assemblage based on the most highly expressed transcripts, which included toxin candidates expressed at >100 CPM among *E. quadricolor* and *C. gigantea* in at least one time point across all samples. Relatively few toxin candidates for *R. crispa*, *S. haddoni*, and *S. helianthus* were expressed at these levels and we reduced the threshold for further evaluation to >10 TPM at a single time point for these species.

### Candidate toxin expression and differential gene expression analysis for the clonal E. quadricolor across the first 48 hours of hosting

Among hosting sea anemones the most abundant toxins remained consistent across all time points, with just two toxin functional groups, putative SCRiPs and the alpha-2-macroglobulin venom factor ( Figure 1), which are not functionally characterized thoroughly in sea anemones. Among the most highly expressed individual transcripts, we recovered toxin candidates corresponded with a poorly characterized toxin candidate previously identified in corals (XP_015758456.1, XP_074617199.1, XP_073247423.1), but also a sea anemone toxin belonging to the SCRiP toxin gene family (A0A3P8MJV5). When comparing AM to PM sampling at the 48 hour mark there were notable differences among the average CPM values observed in the control sea anemones and to a lesser extent among the hosting sea anemones. A single candidate SCRiP gene (C0H693) at the PM 48-hour mark within the control was the most abundant toxin across all individuals and time points. This was further emphasized when evaluating the comparative analysis of TMM values, which allowed sample-wide comparative analysis with normalized values. TMM values for the SCRiP candidates (A0A3P8MJV5, C0H6990, and C0H693) were very high in two individual sea anemones (1 and 2) in the control group, expressed much lower in the AM and almost not at all during the first 48 hours of the hosting groups (Figure 2).

**Figure 1.**
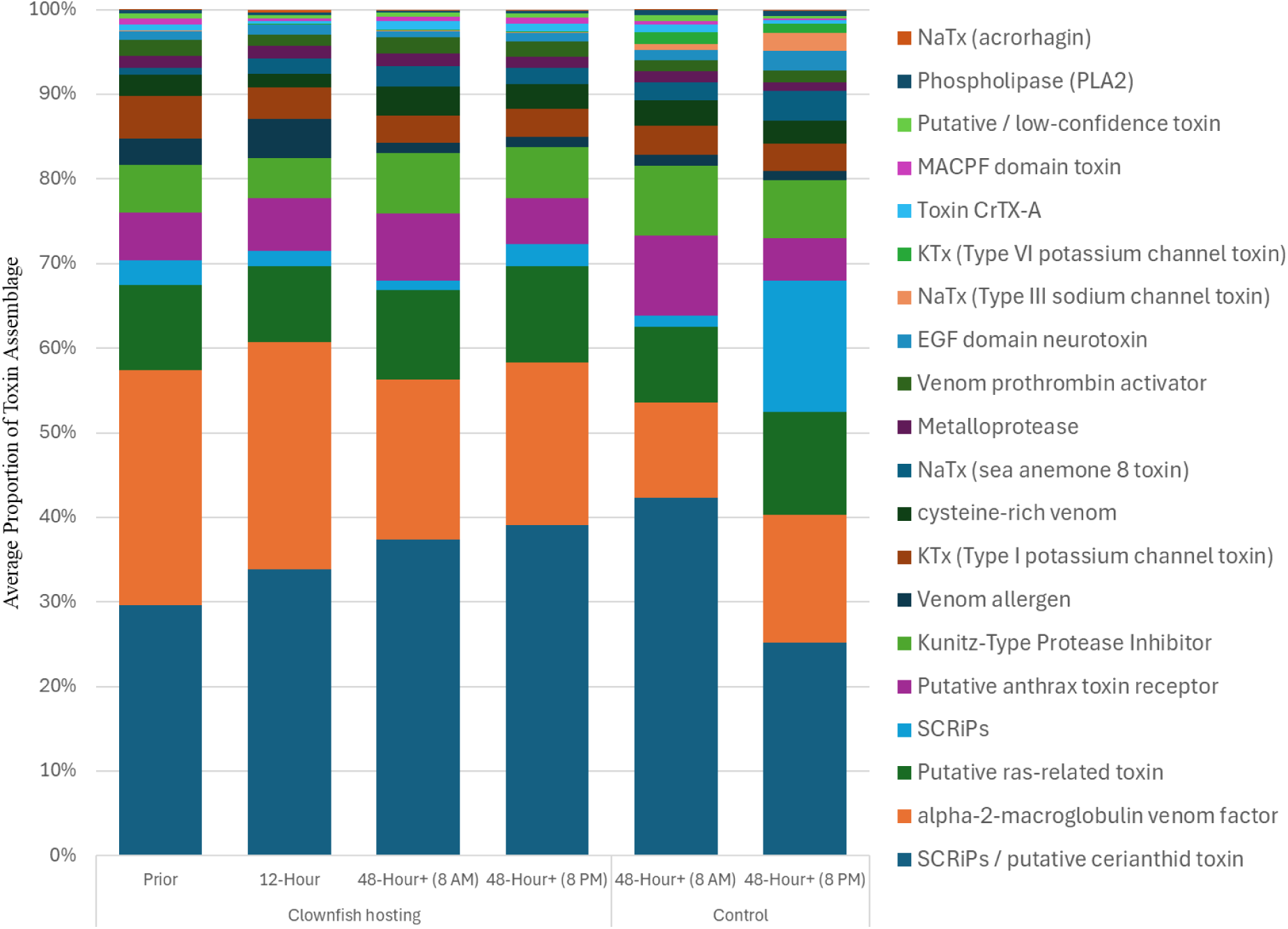
Average cumulative toxin expression (CPM) among hosting *E. quadricolor* sea anemones across the first 48 hours, with controls providing contrast for the AM vs. PM expression profiles. Toxin candidate transcripts with CPM values >100 were combined into distinct functional toxin groups and sorted in the figure legend with the most abundant groups at the bottom of the legend, moving upward as abundances decrease.

**Figure 2.**
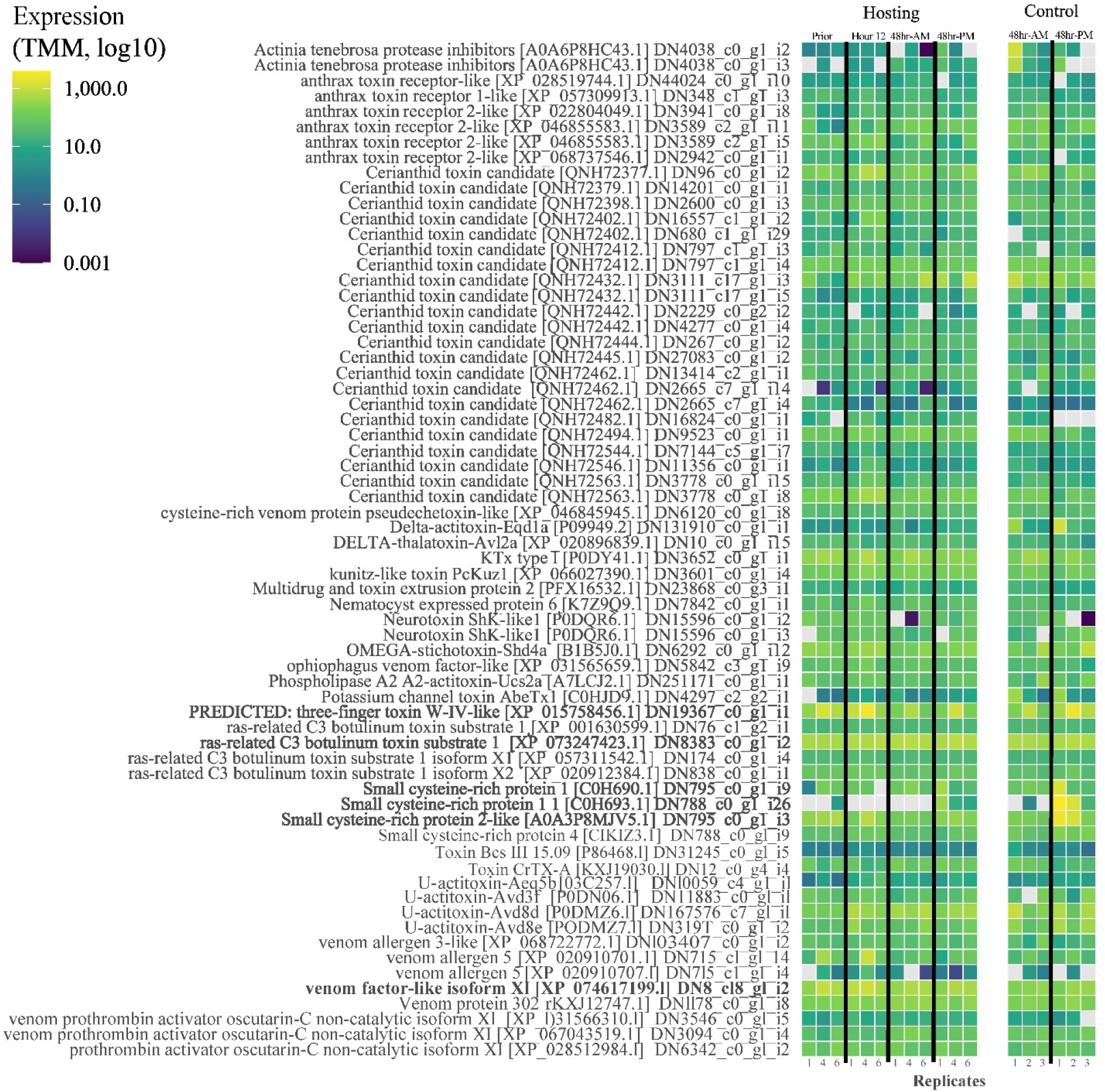
Expression levels as normalized TMM values across all replicates, timepoints, and associations, as well as controls, in *E. quadrioclor*. This heat map only includes toxin-like candidate transcripts with a CPM value >100 for at least one of the biological replicates. Bold toxin candidates are mentioned in the text. Gray cells represent TMM values of zero.

The TMM analysis also allowed us to identify variation among individual contributions to the overall gene expression levels within the hosting group. Specifically, among the most highly expressed transcripts individual number 4 consistently had transcripts with much higher toxin expression levels compared to other hosting sea anemones. Despite the individual variation among some of the more highly expressed transcripts, we observed relatively stable toxin gene expression profiles across all individuals and all time points. There were no drastic changes resulting in dynamic shifts among toxin profiles at any given time point (Figure 2). Furthermore, the differential gene expression analysis only identified one toxin candidate where gene expression profiles demonstrated a significant difference in expression, the SCRiP toxin among 48-hour PM control groups when compared to all other time points, regardless of hosting or control. As our focal *E. quadricolor* was a known clonal propagator, our expanded survey of toxin variation among individuals using previously published data. When looking at the cumulative toxin gene expression levels among individuals they exhibit much more variation compared to Among the individuals, with a single transcript containing a membrane-attack complex/perforin (MACPF) toxin domain (Toxin_AvTX-60A) comprising >70% of the cumulative toxin profile for a single individual, while being completely lost in another (Supplemental Figure 1). The single-copy MACPF toxin dominated five of the six individuals surveyed, with the sixth dominated by a single cytolytic actinoporin. Although the conditions of these individuals remain unknown, it demonstrates that variation in toxin gene is highly likely.

### Toxin expression and transcript shifts for the atypical hosting C. gigantea

Similar to the CPM profiles observed in *E. quadricolor*, the average proportion of each toxin group remained relatively consistent as they contributed to the cumulative toxin expression profiles. Although *C. gigantea* did not differ substantially across sampling time points, there were three toxin functional groups that made up ∼50% of each toxin assemblage, including alpha-2-macroglobulins, actinoporins, and putative cerianthid SCRiPs (Figure 3). Among these, pore forming actinoporins are the only toxin group that has been functionally characterized in sea anemones, with much less known about the alpha-2-macroglobulins and cerianthid SCRiPs. Among the more highly expressed individual transcripts, there was a single sea anemone 8 (SA8) toxin-like (P0DMZ3) gene and two pore-forming actinoporin (Q93109 and C5NSL2) genes (Figure 4). Unlike the *E. quadricolor* anemones, *C. gigantea* were not known to be clonal and all demonstrated slight color morph differentiation, which has been attributed to genotypic variation (Stoletzki and Schierwater, 2005). Although we do not know the origin or environmental conditions of the individuals from the previously published data (BioProject PRJEB21970), these individuals also had much higher variation among their cumulative TPM profiles. Two individuals were dominated by actinoporins (Cytolysin RTX-S-2) at TPM values >30%, with the four others having a combination of four or more toxins contributing to the cumulative toxin profile, with no single toxin consistently contributing more than 25% of the cumulative profile (Supplemental Figure 2).

**Figure 3.**
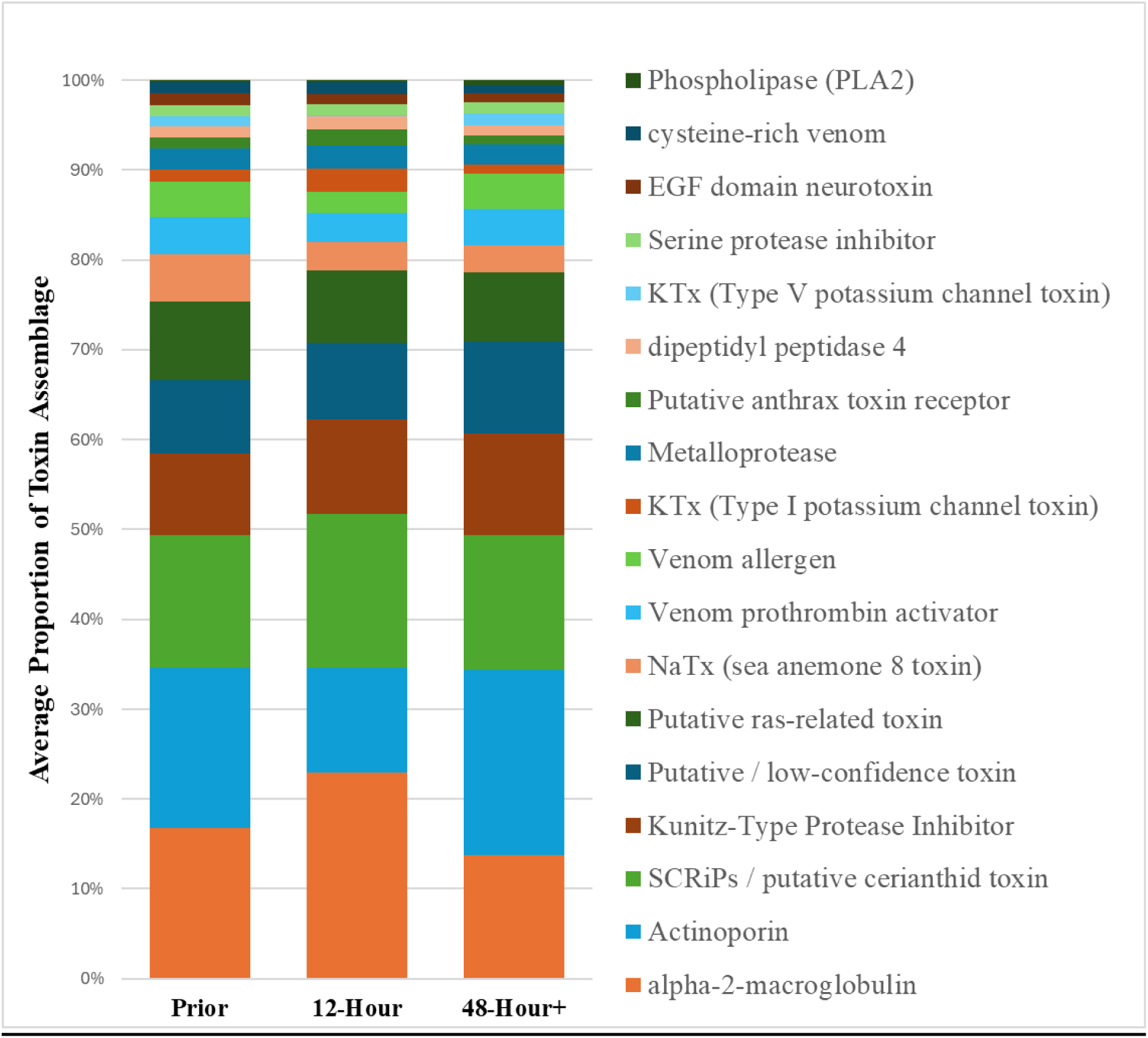
Average cumulative toxin expression (CPM) among atypical hosting *C. gigantea* sea anemones across the first 48 hours. Toxin candidate transcripts with CPM values >100 were combined into distinct functional toxin groups and sorted with the most abundant groups at the bottom of the legend, moving upward as abundances decrease.

**Figure 4.**
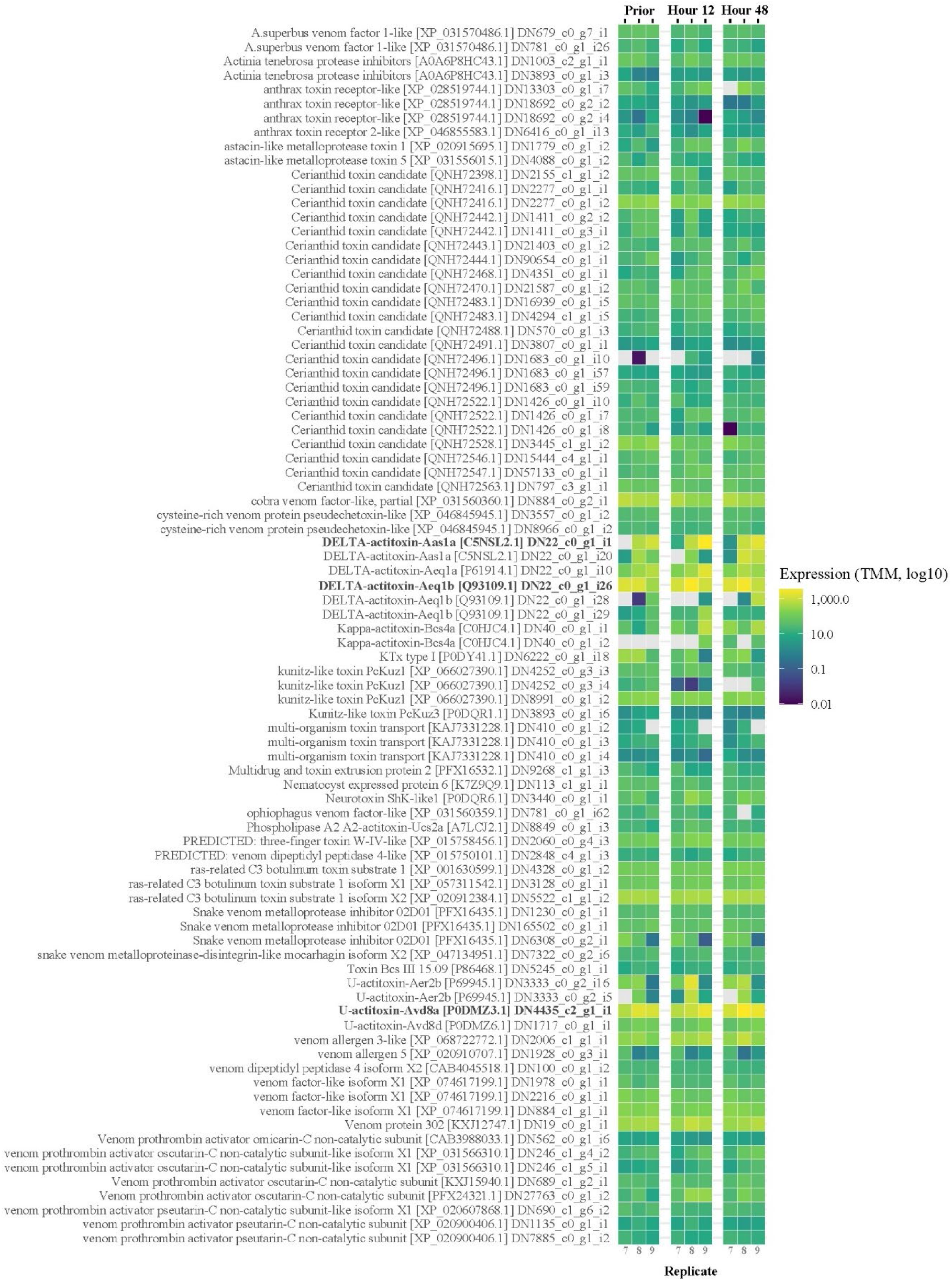
Expression levels as normalized TMM values across all replicates, timepoints, and associations for *C. gigantea*. This heat map only includes toxin-like candidate transcripts with a CPM value >100 for at least one of the biological replicates. Bold toxin candidates are mentioned in the text. Gray cells represent TMM values of zero.

In contrast to *E. quadricolor* TagSeq analysis, we did not have any upregulated MACPF toxins. Instead, the most highly expressed toxin candidates across all time points consisted of nine actinoporins (sequence similarity to C0HJC4, P69928, Q93109, P61914, and C5NSL2), and to a lesser extent the SA8 toxin (P0DMZ3) (Figure 4). The two most upregulated actinoporins did not vary significantly over time following clownfish associations, however, four of the actinoporins with lower expression levels increased following the establishment of a clownfish symbiont (i29, i6, i4, and i10), while three of them decreased (i7, i28, and i20) (Supplemental Figure 3).

### Combined samples and RNA-Seq putative toxin expression levels for R. crispa, S. haddoni, and S. helianthus

In contrast to the TagSeq approach and CPM values for *E. quadricolor* and *C. gigantea*, the TPM values quantified using RNA-Seq for toxin candidates were consistently lower for *R. crispa*, *S. haddoni*, and *S. helianthus*, with only 12, 5, and 4 toxin candidate transcripts expressed at TPM levels >100, respectively. As TagSeq is more sensitive to transcripts exhibiting low to moderate abundance (Lohman et al., 2016), our focus on the more highly expressed transcripts aimed to minimize interpretations or analyses among lowly expressed transcripts that would be misconstrued because of these alternative sequencing approaches. Additionally, at the time of our experimental design and sequencing, the publicly available transcriptomes for *S. haddoni*, and *S. helianthus* were not published and we therefore opted to pursue RNA-Seq at the cost of fewer raw reads involved with mapping. Ultimately, we would have had to sequence 6x the number of raw reads for RNA-Seq to obtain equivalent expression levels to those of TagSeq (Weng and Juenger, 2022), but were only able to sequence 2-3x the number of raw reads. Furthermore, the inability to sequence multiple individuals for biological replicates resulted in our combining three individuals from each time point, which likely resulted in a normalization-like RNA expression profile (Takele Assefa et al., 2020), reducing the frequency of a single individual at a single time point expressing a toxin at high levels at any time point during the experiment. Therefore, for these species, we expanded our TPM threshold to >10 TPM to investigate time point specific shifts in toxin abundance, with a focus on the most highly expressed transcripts.

The cumulative TPM values for *R. crispa* revealed that SA8 toxin and actinoporins comprised the majority of the toxin functional groups, with a proportional increase in metalloproteases at the 12-hour time point (Figure 5). Among these functional groups, there were four toxin candidates contributing >40% of the cumulative toxin gene expression profile, comprising of a cytolytic actinoporin (Q86FQ0), two SA8 toxin-like transcripts (P0DMZ3), and a metalloproteinase that was identified in the *Actinia tenebrosa* genome annotation (XP_031573922). Notably among the 12 toxin-like transcripts at TPM levels >100, five transcripts shared high sequence similarity to the SA8 toxin (Supplemental File 1). Among the TMM analysis subtle shifts across time become more apparent when including multiple toxins at TPM values >10. The cytolytic actinoporins shared high sequence similarity with sagatoxin (Q86FQ0), while other actinoporins (Q93109) were expressed at comparatively lower levels (Figure 6). The BBH toxin transcript (A0A0P0UTI6), which was recently characterized in *Heteractis aurora* and shown to be potent to crustaceans (Homma et al., 2024), was upregulated at the 12-hour time point relative to the prior and 48-hour samples (Figure 6). Apart from the SA8 toxins, all other toxin groups remain relatively stable or expressed at low levels across all time points.

**Figure 5.**
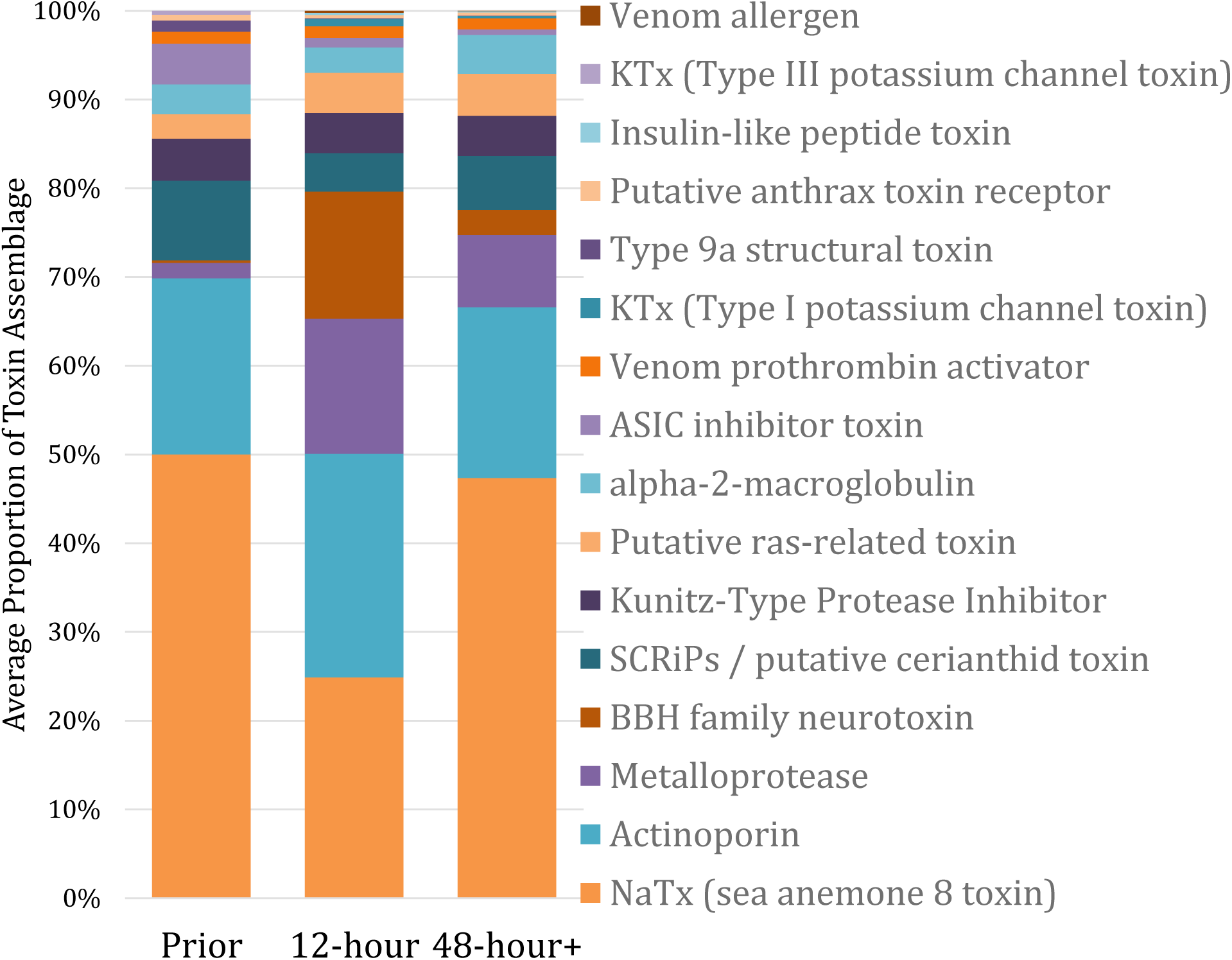
Average cumulative toxin expression (TPM) among *R. crispa* across the first 48 hours. Toxin candidate transcripts with TPM values >10 were combined into distinct functional toxin groups and sorted with the most abundant groups at the bottom of the legend, moving upward as abundances decrease.

**Figure 6.**
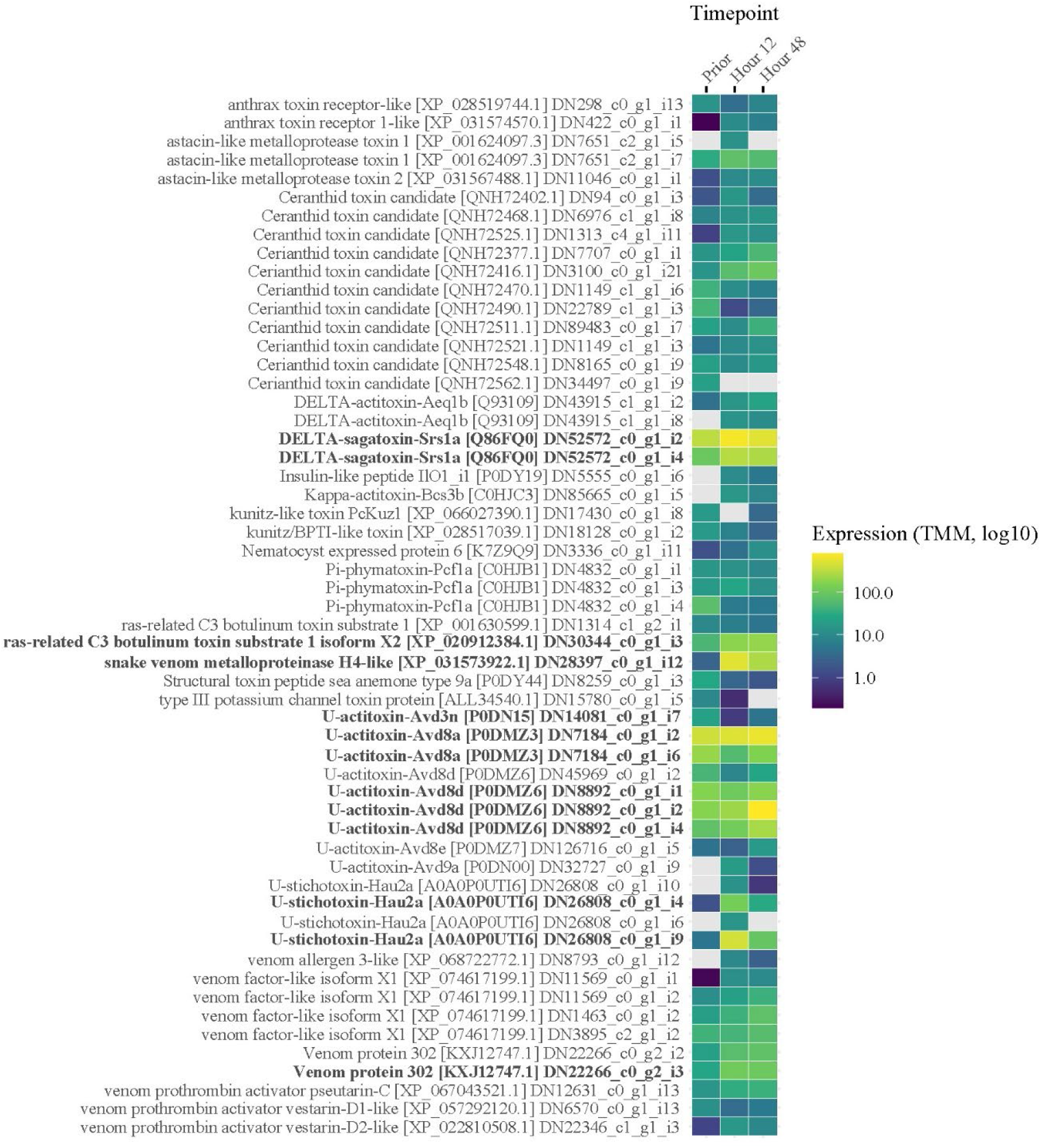
Expression levels as normalized TMM values across all replicates, timepoints, and associations for *R. crispa*. This heat map only includes toxin-like candidate transcripts with a TPM value >10 for one of the three samples. Bold toxin candidates are mentioned in the text. Gray cells represent TMM values of zero.

Among our *S. haddoni* toxin candidates, we observed high expression of Kunitz-Type Protease inhibitors and the SA8 toxin-like groups, with a similar shift at the 12-hour time point with the putative cerianthid SCRiPs and Type III potassium channel toxins increasing their relative proportion (Figure 7). A single SA8 toxin-like (P0DMZ6) gene dominated the cumulative transcriptome, ranging from 20 −40% of the total contribution to the recovered toxin assemblage (Supplemental file). Prior to clownfish association it was the only toxin transcript expressed at >100 TPM with the other four toxin expression levels, which transcripts all shared high sequence similarity to protease inhibitors (B2G331, C0HJU6, C0HLS4, B2G331), increased only after 48 hours of clownfish associations (Figure 8). Beyond the protease inhibitors, there were no single toxin groups or transcripts that seemed to depict any trends correlating with time spent establishing symbiotic associations.

**Figure 7.**
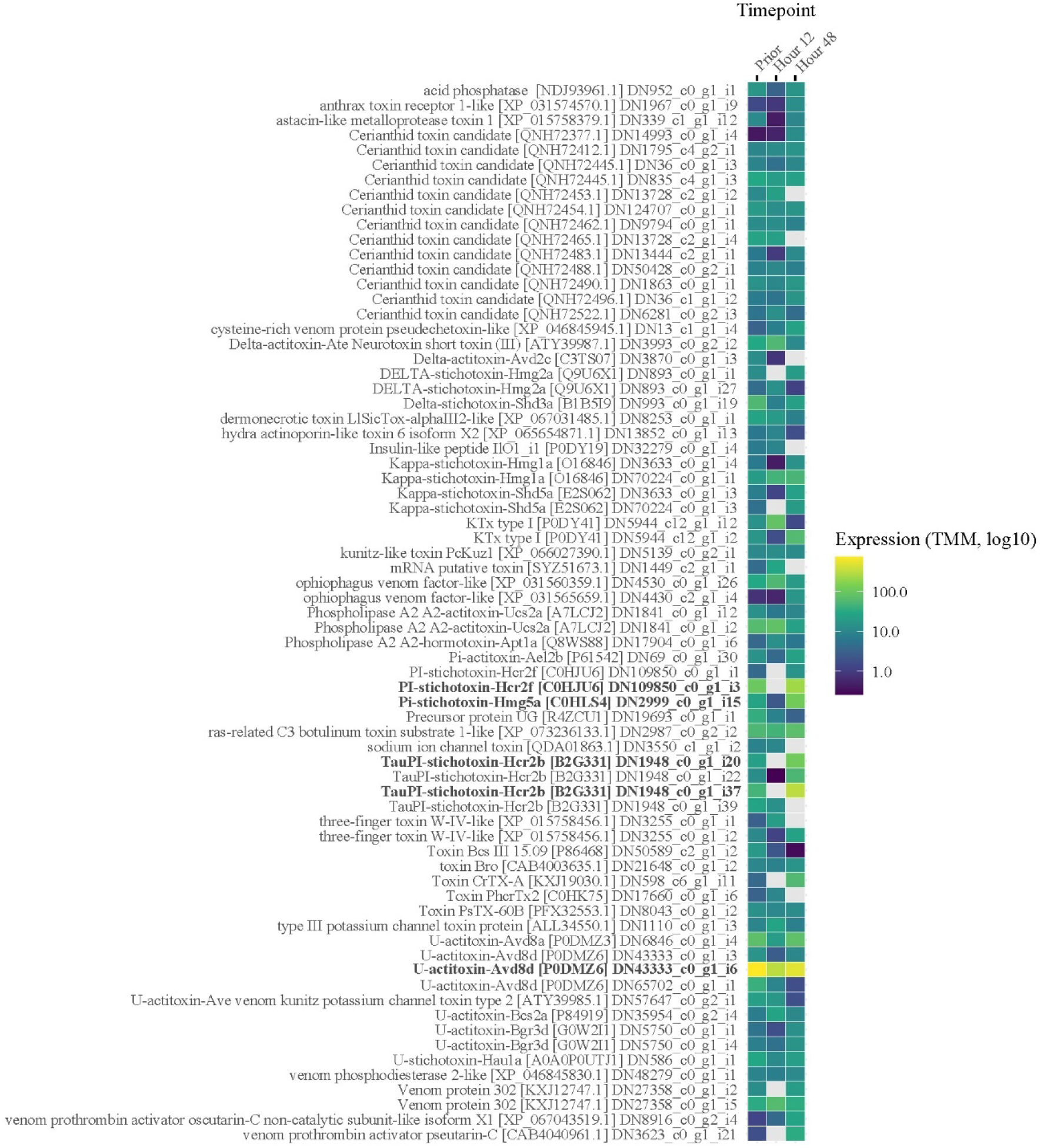
Expression levels as normalized TMM values across all replicates, timepoints, and associations for *S. haddoni*. This heat map only includes toxin-like candidate transcripts with a TPM value >10 for one of the three samples. Bold toxin candidates are mentioned in the text. Gray cells represent TMM values of zero.

**Figure 8.**
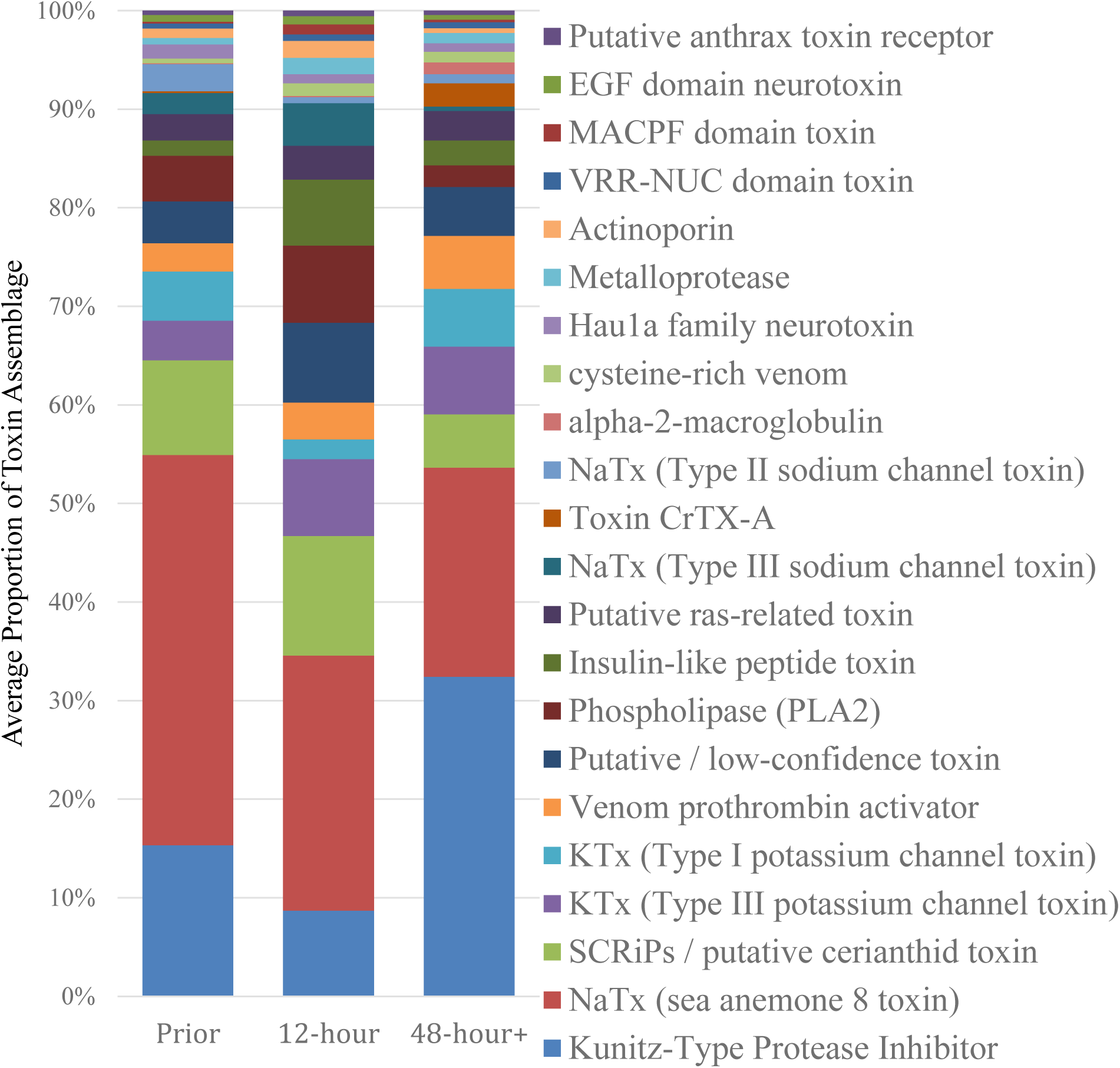
Average cumulative toxin expression (TPM) among *S. haddoni* across the first 48 hours. Toxin candidate transcripts with TPM values >10 were combined into distinct functional toxin groups and sorted with the most abundant groups at the bottom of the legend, moving upward as abundances decrease.

Ultimately for *S. helianthus* the cumulative toxin profiles comprised of SA8 toxins and the poorly characterized putative ras-related toxin groups (Figure 9). Although labeled as “functional groups” these were each dominated by single transcript, a single SA8 toxin-like (P0DMZ6) and a ras-related C3 botulinum toxin (XP_028395254.1) gene (Figure 10). The ras-related C3 botulinum toxin is likely not an actual sea anemone toxin, as it was simply identified in the *Dendronephthya gigantea* (Octocorallia) genome annotation pipeline, we did not remove any unlikely toxin candidates from our results to keep the results consistent through this study. There was notably much less variation among the S. helianthus TMM values across sampling time points indicating that toxin expression did not vary significantly during initial symbiotic association (Figure 10).

**Figure 9.**
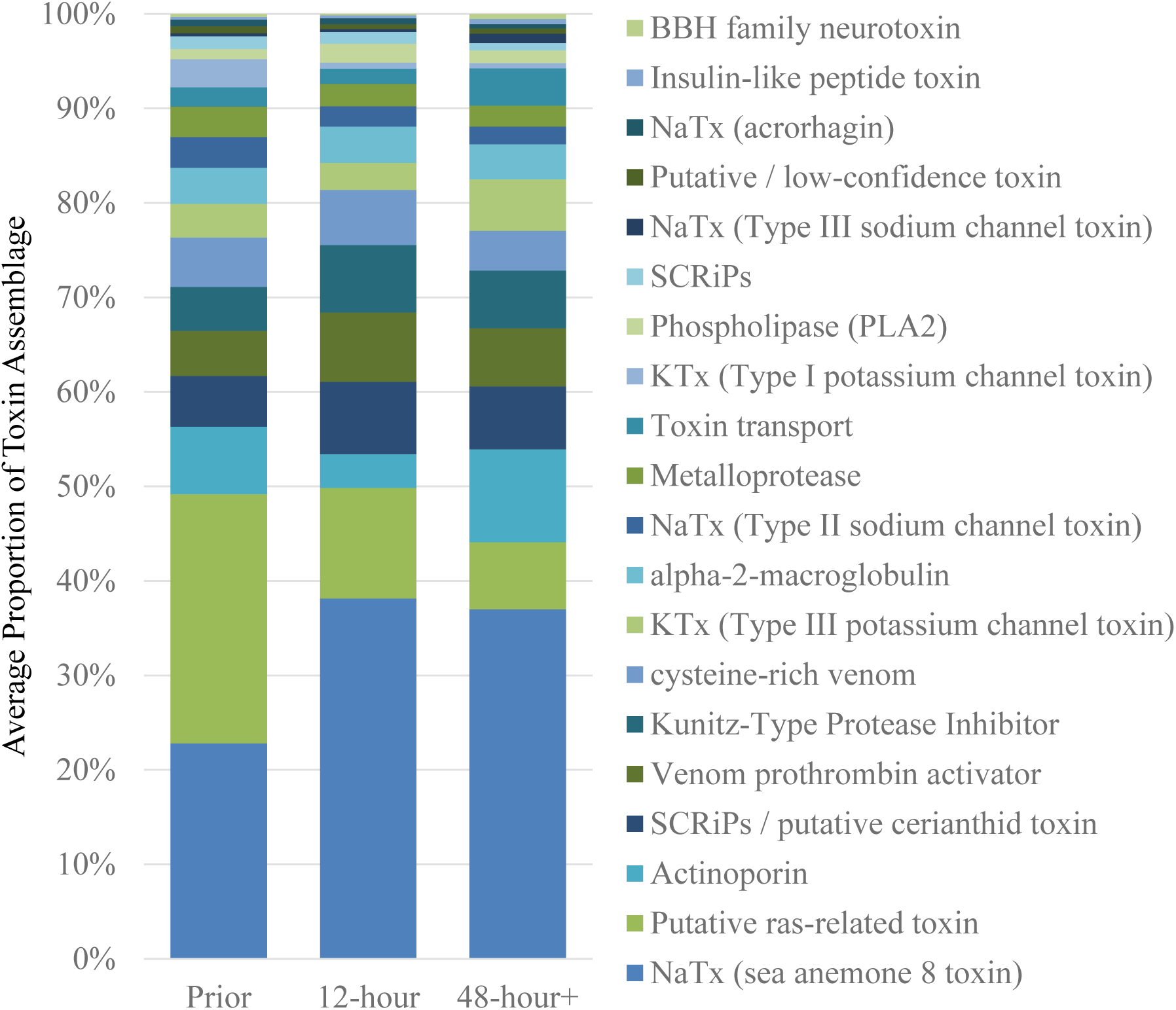
Average cumulative toxin expression (TPM) among *S. helianthus* across the first 48 hours. Toxin candidate transcripts with TPM values >10 were combined into distinct functional toxin groups and sorted with the most abundant groups at the bottom of the legend, moving upward as abundances decrease.

**Figure 10.**
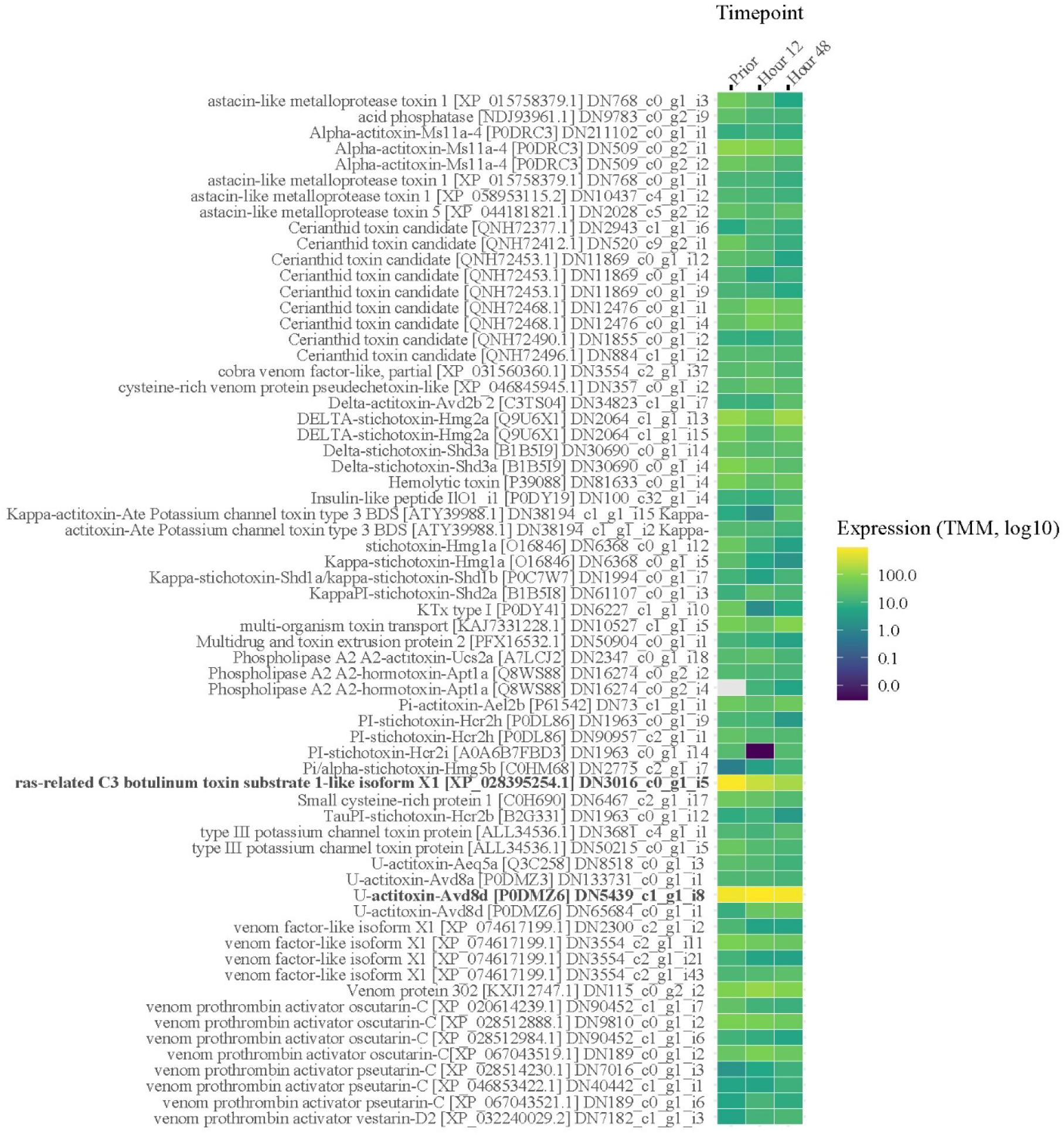
Expression levels as normalized TMM values across all replicates, timepoints, and associations for *S. helianthus*. This heat map only includes toxin-like candidate transcripts with a TPM value >10 for one of the three samples. Bold toxin candidates mentioned in the text. Gray cells represent TMM values of zero.

## Discussion

Our results demonstrate that the initial association between sea anemones and their clownfish symbiont do not significantly influence overall toxin gene expression in typical or atypical hosting sea anemones. Although the life history and physiological processes involved with shaping this mutualistic symbiosis has been well studied (Holbrook and Schmitt, 2005; Porat and Chadwick-Furman, 2005, 2004; Roopin et al., 2008; Schmiege et al., 2017), to our knowledge this is the first investigation into transcriptomic changes associated with establishing this symbiosis. Subtle changes or shifts in toxin gene expression were observed among some candidate toxins (i.e. Cytolytic actinoporin transcript variant i28 within *C. gigantea*, Supplemental Figure 3), however this transcript was not significantly different in its expression profile across time points according to our differential gene expression analysis. Across both *E. quadricolor* and *C. gigantea* only a single toxin transcript was shown to be differentially expressed. However, this transcript corresponded to the 48-hour PM control group when compared to all other groups. The significantly different toxin-like transcript corresponded to a small cysteine-rich protein (SCRiP) (C0H693), but its upregulation during a PM control group vs. AM is most likely due to circadian or other cues rather than anemonefish hosting (Leach et al., 2018; Oren et al., 2015). We did see several toxin candidates among *R. crispa*, *S. haddoni*, and *S. helianthus* decrease after association was established, but even when considering a 4-fold drop among TMM values it was observed in approximately 40% of the transcripts in *S. haddoni* and in *R. crispa* and *S. helianthus* it was observed in 13% and 14% of the transcripts, respectively. Conversely, if we would expect venom expression to increase after association to protect the clownfish we see a 4-fold increase in *R. crispa* in approximately 40% of the transcripts, but to a lesser extent in *S. haddoni* (21%) and *S. helianthus* (5%). Therefore, these general trends indicate that shortly after symbiosis establishment clownfish sea anemones do not decrease overall toxin assemblage to accommodate clownfish symbionts, nor do they increase their overall toxin profile to protect their mutualistic partners against predatory conspecifics.

Across four of the five species surveyed SA8 toxins were consistently among the most highly abundant toxin-like transcripts recovered in our analysis. The role of SA8 toxins broadly across sea anemones are poorly understood, however, a recent investigation by Ashwood et al. (2023) was able to comprehensively investigate this elusive toxin. Although the SA8 toxins were originally identified when characterizing the EST library for *Anemonia viridis* (Kozlov and Grishin, 2011). Since then, its disulfide connectivity has been resolved, resembling the ShK motif and found across more than 14 species in various tissues. Genomic infrastructure surrounding this toxin candidate indicates that it sometimes exhibits copy number and arrangement variants and expansions coupled with gene inversions (Ashwood et al., 2023). Although transcripts SA8 toxins were found to be differentially expressed among tissues and recovered within the venom proteome of *Telemactis stephensoni* (Ashwood et al., 2023), in *E. quadricolor* it was not recovered in the proteome despite recovering 10 gene clusters among the transcripts (Hoepner et al., 2024). Processes shaping the genomic infrastructure surrounding the SA8 toxin across sea anemones mirrors that of what was recently described at the population level in *Nematostella vectenssi* (Smith et al., 2023b). Although the SA8 toxin has not been shown exhibit strong paralytic or potassium channel blocking activities (Ashwood et al., 2023),future investigation into toxin gene family evolution within species having high SA8 copy number may reveal divergent functional properties among paralogs, similar to what has been previously reported in *N. vectensis* (Sachkova et al., 2019).

Although preliminary, our results may indicate the downregulation, or shift in gene expression, of a single cytolytic actinoporin transcript variant essential to the establishment of symbiosis in *C. gigantea*. The addition of *C. gigantea* genomic datasets (BioProject: PRJEB78657) and other members across Actinioidea is warranted to thoroughly elucidate evolutionary processes shaping these repertoires at the genomic level. Preliminary hosting testing the ability of *C. gigantea* to establish these associations, individual 9 inadvertently resulted in two clownfish mortalities when trying to establish this association. While we did not identify the precise cause of mortality at the time, the prior tentacle condition had a single actinoporin (i28 from DN22_c0_g1) upregulated at almost 600x higher when compared to later time points, as well as the other two *C. gigantea* sea anemones (Figure 4). When aligned, this actinoporin differed at two key points from other at only two amino acid resides (Q65P and N69H) from other known actinoporin gene copies found within *C. gigantea* (Supplemental Figure 3) and more broadly across sea anemones (Macrander and Daly, 2016). Aside from these two unique amino acid substitutions, they were almost identical to previously published *C. gigantea* and other sea anemone actinoporin sequences analyzed within Macrander and Daly (2016), with no noteworthy changes in the previously published actinoporin gene tree reconstruction (data not shown). The site-specific mutations of these potential toxins have not been characterized previously, but were found adjacent to the oligomerization site, which may play a role in pore formation (Madio et al. 2019; Anderluh and Maček 2002). Although this is not conclusive evidence of a barrier to atypical sea anemone hosts in establishing clownfish symbioses, there could be something specific within the single cytolytic actinoporin gene preventing symbiont association. It is not unreasonable to predict that a shift in toxin abundance would be a necessary precuring to permit symbiosis, as clownfish hosting sea anemones has evolved independently three different times (Kashimoto et al., 2022; Titus et al., 2019). Variation in toxin assemblage across this study indicates that expression likely fluctuates, despite having a consistent overall repertoire to build their toxin assemblage.

Our resampling of the same individuals is likely a key reason behind the stable toxin assemblage observed over time when symbiont hosting is established. Individual variation in cumulative toxin expression for the publicly available *E. quadricolor* (Supplemental Figure 1) and *C. gigantea* (Supplemental Figure 2) indicated that although there are for the most part consistent toxin expression profiles across individuals, toxin expression profiles can change drastically among individual sea anemones. Many of the sea anemone toxin studies thus far have either combined individual samples or sampled a single individual, which could have significant implications on our interpretation of species-specific toxin assemblages. Furthermore, the short exposure time was selected due to previous investigations into *N. vectensis* respiration following venom discharge correlated with the upregulation of key venom components after just 2 hours (Sachkova et al., 2020). Because of this we anticipated that once symbiosis was established, if toxins changed their expression profiles accordingly we would see toxin gene expression shifts occurring within the first 12 hours. However, the ability for clownfish to hide amongst the tentacles likely minimizes the frequency at which venom is discharged when this symbiosis is established (Hoepner et al., 2022; Mebs, 1994). Instead, long-term association studies may be better suited to accurately explore the physiological relationship between clownfish hosting dynamics and shifts among key life history traits that have shown to be benefits of this symbiosis (Holbrook and Schmitt, 2005; Porat and Chadwick-Furman, 2005, 2004; Schmiege et al., 2017).

While our primary analysis remains focused on the toxin repertoire and expression dynamics, we did recover 715 DEGs in *E. quadricolor* and 251 in *C. gigantea* that do not encode for toxins. Although a full functional characterization is beyond the scope of this current manuscript, preliminary tBLASTn searches (Altschul et al., 1990) against the UniProt dataset (The UniProt Consortium, 2019) provided insight into their putative function. Many of the more highly recovered gene ontology groups linked to metabolic pathways, nutrient transport, and peroxidase activities, underscoring the broad physiological shifts triggered by symbiosis establishment. Given these preliminary findings, we are planning to expand our analysis to further evaluate the gene expression profiles of the sea anemones and their zooxanthellae symbionts, which are all essential components of these beneficial mutualisms. Furthe exploration into symbiotic gene expression dynamics beyond the venom system will likely identify pathways facilitating nutrient exchange and metabolic integration, even among clonal individuals with low genetic variation. A comprehensive investigation into these non-toxin pathways, including a broader comparative ‘omics analysis of metabolism and symbiont gene repertoires, will be the subject of a forthcoming paper, as the complexity of these metabolic shifts warrants a dedicated and detailed exploration.

We acknowledge that the prior experience or ‘naivety’ of the clownfish represents an uncontrolled variable in this study. While our results show stable toxin profiles regardless of the fish’s history, the potential for host experience to subtly influence anemone gene expression cannot be statistically ruled out with the current level of replication. Future studies with higher replicate counts per species would be better equipped to model the interactions between symbiont naivety and host response. Future comparative gene expression studies which aim to elucidate molecular processes surrounding clownfish hosting establishment and its maintenance should use both short and long-term sampling, as well as sufficient replicates to evaluate the role of naivety in clownfish. These approaches would more accurately quantify molecular mechanisms and responsible for establishing and maintaining this broadly familiar and charismatically popular symbiotic relationship.

## Supporting information

Supplemental File- CPM/TPM/TMM values

## Acknowledgements

The authors would like to thank Kerry Broderick for her support with animal care and data analysis. The authors would like to thank Ashley Bowers-Macrander for contributing significant effort to support in the data analysis and writing of this manuscript. We would also like to thank Darrell’s Aquarium Store, and Caribbean Tropicals. Several students were supported through the Florida Southern College Hansen Science Scholars Program.

---

**Supplemental Figure 1.**
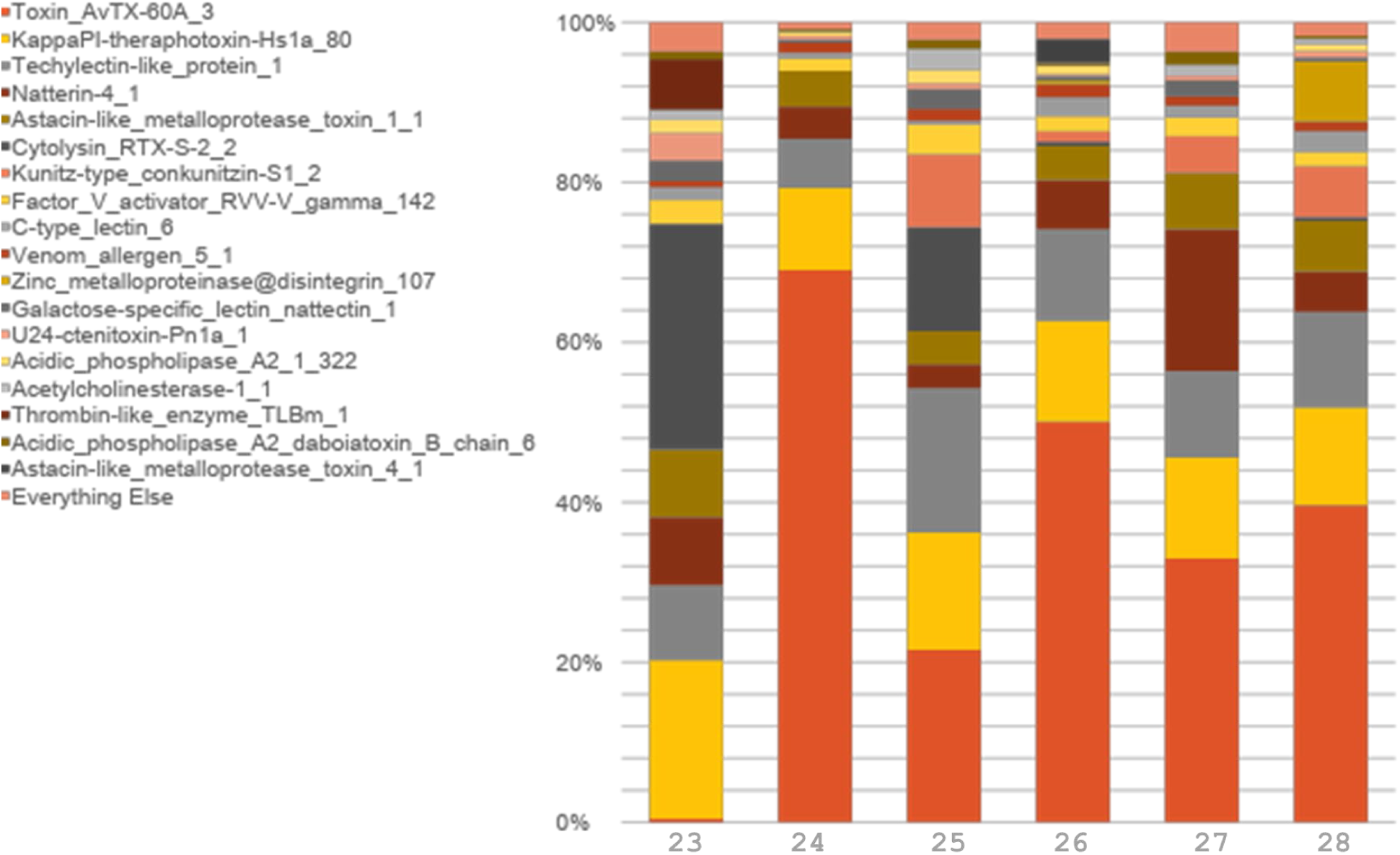
Cumulative expression (TPM) among toxin candidates among six individual *E. quadricolor* from previously published datasets (BioProject PRJEB21970). X-axis labels correspond to the last two digits of the run ID (ERX2104223 -ERX2104228). Toxins are sorted in the figure legend, with most abundant toxins at the top of the legend, moving down as abundances decrease.

**Supplemental Figure 2.**
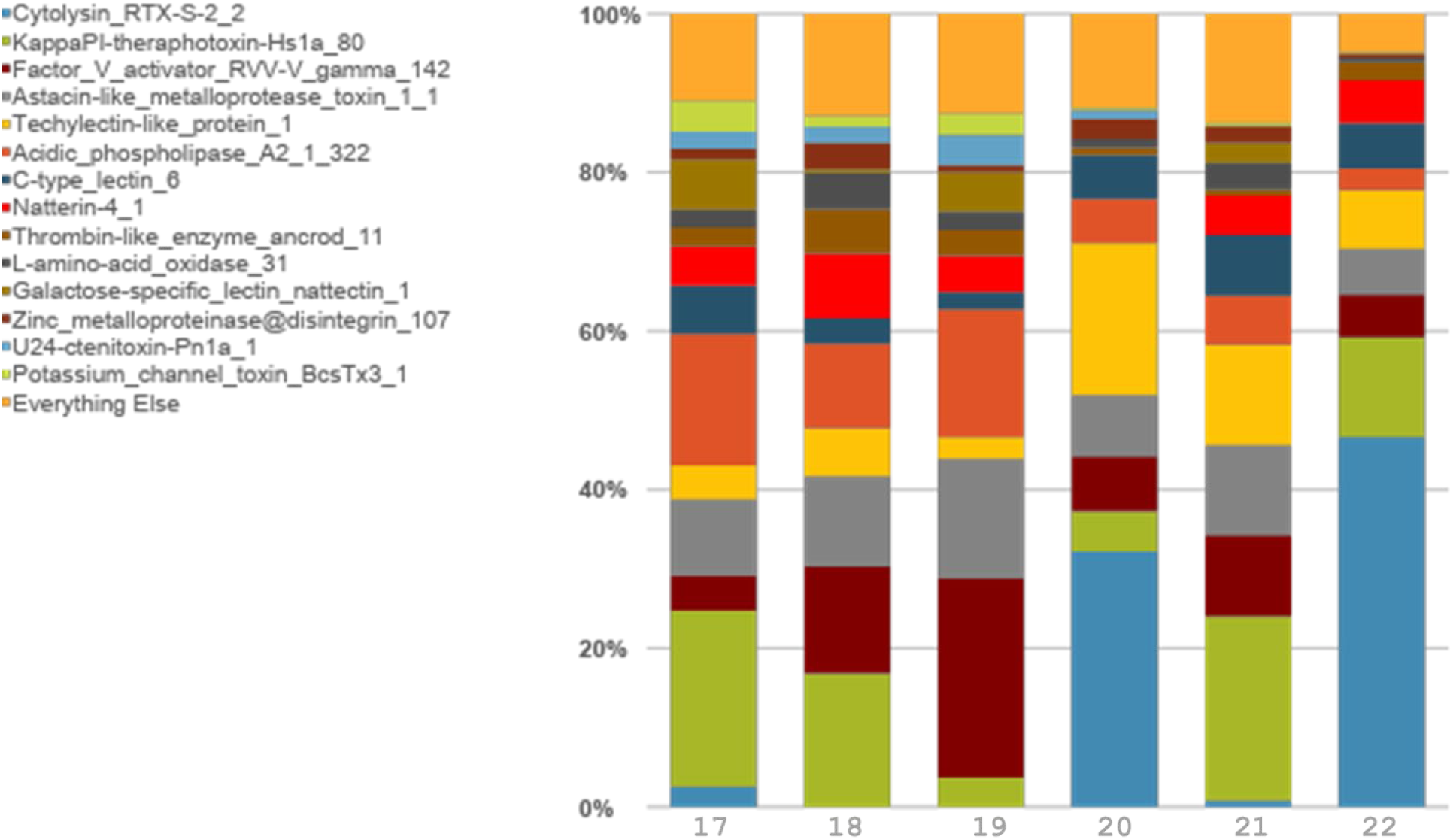
Cumulative expression (TPM) among toxin candidates among six individual *C. gigantea* sea anemones from previously published datasets (BioProject PRJEB21970). X-axis labels correspond to the last two digits of the run ID (ERX2104217 -ERX2104222). Toxins are sorted in the figure legend, with most abundant toxins at the top of the legend, moving down as abundances decrease.

**Supplemental Figure 3.**
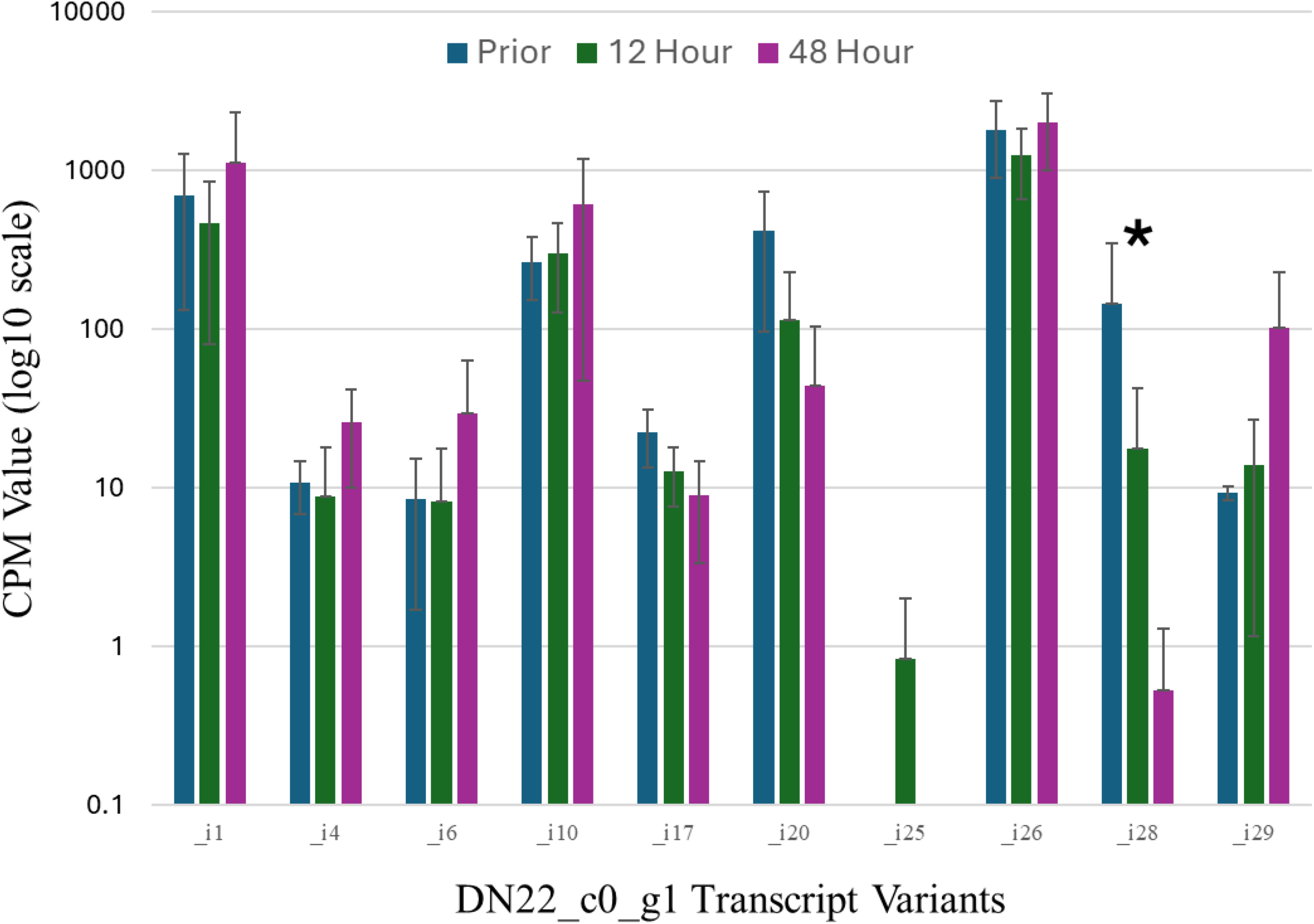
Actinopirin expression (CPM values) across clownfish associations for various putative actinoporin transcript variants. The asterisk (*) highlights a notable fold change corresponding with strong upregulation among individual 9.

**Supplemental Figure 4.**
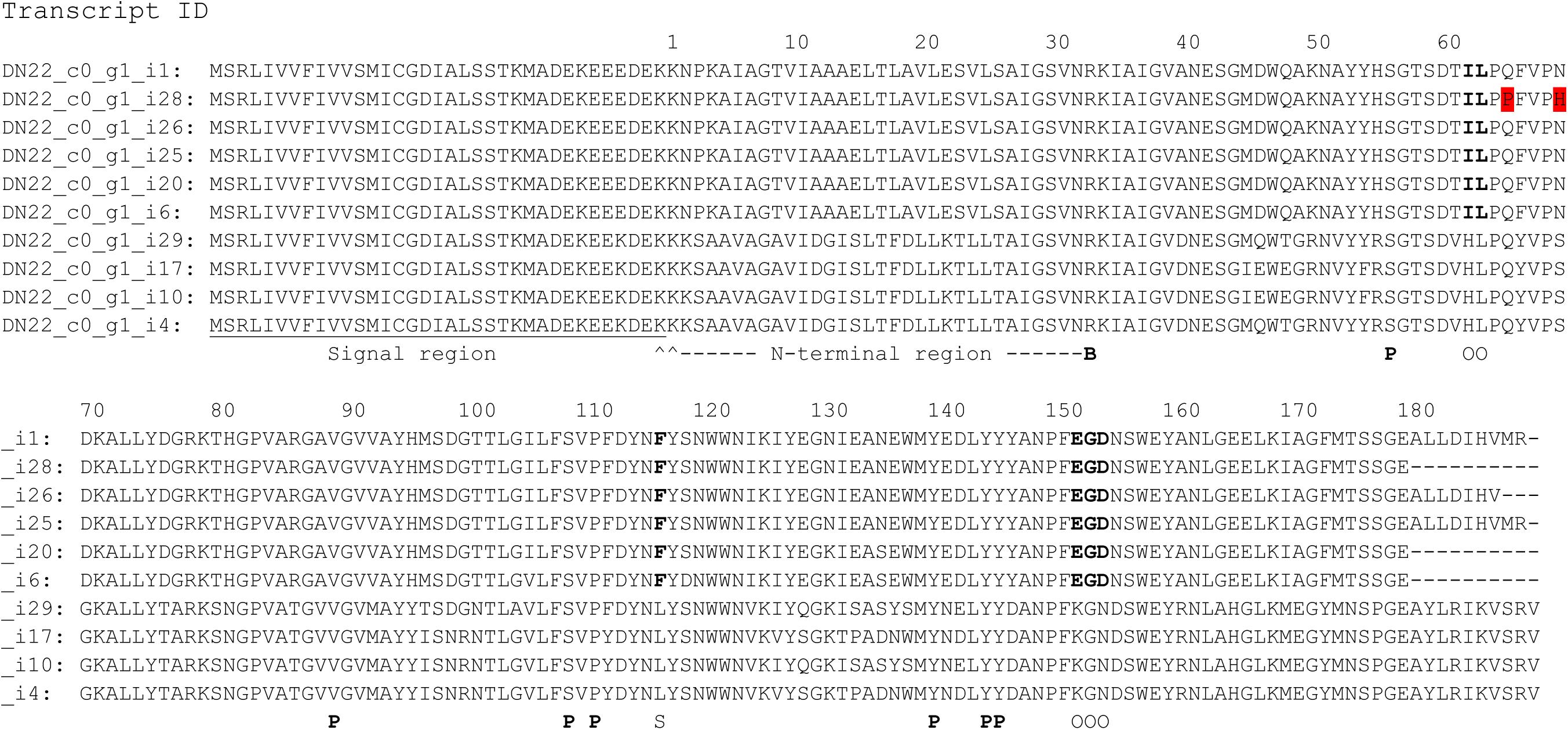
Sequence alignment of DN22_c0_g1 transcript variants. Unique amino acid residues not found in other sea anemone actinoporins adjacent to the oligomerization site are highlighted in red. The signal and N-termianl regions are indicated, with functional sites labeled as (B) site of bend when N-terminus comes into contact with the cell membrane, (P) residues involved with the POC binding site, (O) residues involved with oligomerization, (S) key sphingomyelin binding site.

